# Highly Multiplexed Genome Engineering Using CRISPR/Cas9 gRNA Arrays

**DOI:** 10.1101/331371

**Authors:** Morito Kurata, Natalie K. Wolf, Walker S. Lahr, Madison T. Weg, Samantha Lee, Kai Hui, Masano Shiraiwa, Beau R. Webber, Branden Moriarity

## Abstract

The CRISPR/Cas9 system is an RNA guided nuclease system that evolved as a mechanism of adaptive immunity in bacteria. This system has been adopted for numerous genome engineering applications in research and recently, therapeutics. The CRISPR/Cas9 system has been largely implemented by delivery of Cas9 as protein, RNA, or plasmid along with a chimeric crRNA-tracrRNA guide RNA (gRNA) under the expression of a pol III promoter, such as U6. Using this approach, multiplex genome engineering has been achieved by delivering several U6-gRNA plasmids targeting multiple loci. However, this approach is limiting due to the efficiently of delivering multiple plasmids to a single cell at one time. To augment the capability and accessibility of multiplexed genome engineering, we developed an efficient golden gate based method to assemble gRNAs linked by optimal *Csy4* ribonuclease sequences to deliver up to 10 gRNAs as a single gRNA array transcript. Here we report the optimal expression of our guide RNA array under a strong pol II promoter. This system can be implemented alongside the myriad of CRISPR applications, allowing users to model complex biological processes requiring numerous gRNAs.

## INTRODUCTION

With advances in our understanding of complicated biological processes, such as cancer development and transdifferentiation, the need to readily model and manipulate these complex systems is ever increasing. Being able to modulating multiple genes of the same pathway or separate pathways at one time, will help further our understanding of the functional outcome. The CRISPR (Clustered regularly interspaced short palindromic repeats) and CRISPR associated protein 9 (CRISPR/Cas9) system has become a powerful tool for modifying gene expression. In prokaryotes the CRISPR/Cas9 system functions as an adaptive immune system via targeted disruption of invading phages [1]. In the bacterial genome, the Cas operon consists of variable CRISPR RNA (crRNA) sequences in series with intermittent Csy4 spacer motifs. When transactivated, multiple crRNAs are transcribed in tandem, they are then cleaved into individual pre-crRNAs by Cas6 (also known as Cs) Individual variable pre-crRNA sequences then associate with Cas9 and the invariable tracrRNA, which recruits RNase III to cleave the RNA heteroduplex and form mature crRNA [2]. At the 5’ end of the crRNA is the 20 bp protospacer region which hybridizes with a target DNA sequence and induces a double stranded break (DSB). In synthetic gene editing, the crRNA and tracrRNA can be fused into a single chimeric gRNA that can be cloned into an expression vector without requirements of post-transcriptional modification, where introduction of a gRNA alongside a Cas9 expression vector is a simple and elegant method for targeted DSB induction [1, 2].

CRISPR/Cas9 has now been coopted by numerous laboratories for various genome engineering purposes [2]. This coopted CRISPR/Cas9 system largely utilizes the Cas9 protein along with a single chimeric crRNA-tracrRNA (sgRNA) to localize to target DNA sequences within the genome and induce a DSB. The DSB can be utilized for gene knockout through non-homologous end joining (NHEJ) repair or sequence modification through homology directed repair (HDR) using a donor DNA substrate. Moreover, many groups have utilized the nuclease dead Cas9 (dCas9) mutant for transcriptional activation (VP64, MS2:p65:HSF1, and p300 fusions) [3,4] and repression (KRAB fusion) [5] by directly fusing the heterologous domains to dCas9 or localizing them using the MS2 RNA binding protein in conjunction with a modified gRNA architecture containing MS2 binding sites.

The ability to deliver multiple gRNAs simultaneously makes our CRISPR/Cas9 system highly amenable to multiplex genome editing. Previously, this has been achieved by delivering multiple plasmids encoding gRNAs expressed from the strong U6 pol III promoter targeting several different genes. However, this becomes troublesome due to poor efficiency and toxicity associated with the delivery of large amounts of DNA plasmids to cells. An alternative approach is to deliver the gRNAs as *in vitro* transcribed (IVT) RNAs [6] or chemically synthesized RNA oligonucleotides [7] and Cas9 protein. However, for basic research this approach becomes cost prohibited and is reserved for use in sensitive applications such as gene editing of primary cell types, which do not tolerate transfection with dsDNA/plasmids/expression plasmids. In light of this, a number of groups have generated methods to clone multiple pol III promoter:gRNA elements into a single plasmid in tandem [8] or alternating the orientation array format [9]. Although these methods are minimally functional, there are drawbacks to these approaches. For instance, multiple groups have raised the concern that numerous pol III promoters may engage in “Promoter Cross talk Effects” and reduce the transcriptional activity of endogenous pol III driven genes, potentially causing undesirable side effects [10-12]. Further, as these systems have largely been validated via transient transfection, there may be a large amount of promoter interface when stably integrated, as has been reported using arrays of pol II driven elements [13, 14], potentially limiting their utility for stable CRISPR/Cas9 mediated gene activation or repression. Inspired by the endogenous CRISPR system, herein we describe methods for highly efficient assembly and expression of CRISPR/Cas9 gRNA arrays using golden gate assembly methods and pol II promoters [15]. We validated the system using gRNA array vectors containing up to ten unique gRNAs in human cells through transient expression for gene knockout and activation.

## Material and methods

### Design and construction of guide RNAs

Guide RNAs (gRNAs) were designed to the desired region of each target gene using the CRISPR Design Program (http://crispr.mit.edu). Multiple gRNAs were chosen based on the highest ranked values determined by avoiding off-target locations. The gRNAs were ordered in oligonucleotide pairs, annealed, and ligated into pENTR221-U6 stuffer [16]. Briefly, oligonucleotide pairs were annealed and phosphorylated using T4 PNK (New England BioLabs) and 10X T4 Ligation Buffer (New England BioLabs) in a thermocycler with the following protocol: 37°C 30 minutes, 95°C 5 minutes and then ramped down to 25°C at 5°C/minute. pENTR221-U6 stuffer vector backbone [16] was digested with FastDigest BbsI (Fermentas), FastAP (Fermentas) and 10X Fast Digest Buffer and used for the ligation reaction. The digested pENTR221-U6 stuffer vector was ligated together with the phosphorylated and annealed oligo duplex (1:200 dilution) from the previous step using T4 DNA Ligase and Buffer (New England BioLabs). The ligation was incubated at room temperature for at least 1 hour and then transformed and mini-prepped using the GeneJET Plasmid Miniprep Kit (Thermo Scientific). Plasmids were analyzed by Sanger sequencing to confirm proper oligonucleotide ligation.

### Validation of guide RNAs

HEK293T cells were maintained in DMEM medium supplemented with 10% fetal bovine serum (FBS). 1×10^5^ cells of HEK293T cells were seeded in 24-well plate the day before transfection. Transfection was performed using Lipofectamine 2000 (Thermo Scientific), following the manufactures protocol. 500 ng of pT3.5-CAG-hCas9, 250 ng of pENTR221- U6-gRNA plasmid and 100ng of pGFP-MAX plasmid, to assess transfection efficiency, (Amaxa) were diluted in 75 μl of OptiMEM and 5 μl of Lipofectamine 2000 was diluted in 75 μl of OptiMEM and then the mixtures were combined. The complete mixture was incubated for 15 min before being added to cells in a drop wise fashion. After 16 hours, the media was changed to fresh DMEM medium containing 10% fetal bovine serum. Cells were incubated for 3 days at 37°C and then genomic DNA was collected using the GeneJET Genomic DNA Purification Kit (Thermo Scientific). Activity of the gRNAs was quantified by Surveyor nuclease assay, gel electrophoresis, and densitometry as described [17].

### Vectors construction and LR clonase reactions

Plasmids containing the ccdB gene were maintained in One Shot® ccdB Survival™ 2 T1R bacteria (Invitrogen). All enzymes used in this study were purchased from New England BioLabs. All pENTR1 vectors were derived originally from pENTR221-GUS (Invitrogen). gBlocks (Integrated DNA Technologies) encoding the pGG and pACPT cassettes were ordered and constructed to contain attB1/2 sites. Single BP Clonase reactions were performed using manufacturer’s protocols with Clonase II enzyme mixes (Invitrogen). The pT3.5-CAG-Csy4-T2A-hCas9 and pT3.5-CAG-MS2-p65-HSF1-T2A-eGFP vectors were generated using a combination of restriction enzyme cloning and Gateway cloning. Briefly, the nucleotides encoding attL1-Csy4-T2A with SbfI and BspHI sites was ordered as a gBlock (Integrated DNA Technologies). The DNA fragment of Csy4-T2A was cloned into pENTR221-hCas9 digested with NcoI. pENTR221-Csy4-T2A-Cas9 was then cloned into pT3.5-CAG-DEST via Gateway LR Clonase reaction, following manufacturer’s instructions. A gBlock encoding MS2-p65-HSF1-T2A-eGFP flanked by attB1/2 was used for BP Clonase reaction with pDONR221 to generate pENTR221-MS2-p65-HSF1-T2A-eGFP, which was subsequently used for LR Clonase reaction with pT3.5-CAG-DEST to generate pT3.5-CAG-MS2-p65-HSF1-T2A-eGFP.

### Golden gate assembly of gRNA arrays

Single pGG vectors were generated using the same oligonucleotide ligation approach described above. Primer sequences can be found in **S1 Table**. Arrays were then generated using golden gate assembly. Briefly, 150 ng of each pGG-gRNA plasmid was combined with 150 ng of the appropriate pACPT vector, followed by addition of 1uL of BsaI (New England BioLabs), 1uL of T4 DNA Ligase and Buffer (New England BioLabs) and water to a total of 20 uL. Golden gate assembly was then carried out using the following thermocycling protocol: (37°C 5 minutes, 16°C 10 minutes x10), 50°C 5 minutes, 80°C for 5 minutes and then cooled to 4°C. One microliter of 25 mM ATP and 1 uL of Plasmid Safe (Epicentre) were then added to the reaction and incubated for 1 hour at 37°C. Assembled plasmids were then transformed into TOP10 bacteria and plated on spectinomycin selection plates with X-gal and mini-preps performed on white colonies using the GeneJET Plasmid Miniprep Kit (Life Technologies). The plasmids were then analyzed by Sanger sequencing to confirm proper gRNA array assembly.

### Plasmid transfection in mammalian cells

HEK293T cells were maintained in DMEM medium supplemented with 10% fetal bovine serum (FBS). 1×10^5^ cells of HEK293T cells were seeded in 24-well plate the day before transfection. Transfection was performed using Lipofectamine 2000 (Thermo Scientific), following the manufactures protocol. 500 ng of pT3.5-CAG-Csy4-T2A-hCas9, 250 ng of pENTR221-U6-gRNA or pACPT array plasmid were diluted in 75 μl of OptiMEM and 5 μl of Lipofectamine 2000 was diluted in 75 μl of OptiMEM and then the mixtures were combined. The complete mixture was incubated for 15 min before being added to cells in a drop wise fashion. After 16 hours, the media was changed to fresh DMEM medium containing 10% fetal bovine serum. Cells were incubated for 3 days at 37°C and then genomic DNA was collected using the GeneJET Genomic DNA Purification Kit (Thermo Scientific). Activity of the gRNAs was quantified by a Surveyor nuclease digest, gel electrophoresis, and densitometry as described [16]. For gene activation experiments, 250 ng of pT3.5-CAG- Csy4-T2A-dCas9-VP64, 250 ng of pT3.5-CAG-MS2-p65-HSF1-2A-eGFP and 250 ng of pENTR221-U6-gRNA or pACPT array plasmid were transfected as above. Cells were incubated for 3 days at 37°C and then RNA was extracted using PureLink® RNA Mini Kit (Thermo Fisher Scientific) and then reverse-transcribed by Transcriptor First Strand cDNA Synthesis Kit (Roche).

### Surveyor nuclease assay

Surveyor assays were performed as previously descried [17]. Briefly, after electroporation of CRISPR/Cas9 plasmids and incubation for 3 days genomic DNA was extracted using GeneJET Genomic DNA Purification Kit (Thermo Scientific), following manufacturer’s instructions. PCR amplicons were generated spanning the Cas9 binding site using Accuprime™ Taq HF (Invitrogen) using the following PCR cycle: initial denaturation at 95°C for 5 min; 40x (95°C for 30 sec, 55°C or 60°C for 30 sec, 68°C for 40 sec); final extension at 68°C for 2 min. PCR amplicons were denatures and annealed as follows: 95°C for 5 minutes, 95-85°C at −2°C/s, 85-25°C at −0.1°C/s, 4°C hold. Primer sequences can be found in **S1 Table**. Three microliters of the annealed amplicon was then diluted with 6 μL of 1x Accuprime PCR buffer and treated with 1 μL of Surveyor nuclease with 1 μL of enhancer (Thermo Scientific) at 42°C for 20 min. The reaction was then stopped by the addition of 3 μL of 15% Ficol-400 and 0.05% Orange G solution containing 1 mM EDTA and subsequently run on a standard 10% TBE gel. Percent gene modification was calculated using Image J software as described [2].

### Q-RT-PCR analysis

Taq-man quantitative PCR was performed with following primer and probes; ASCL1; Hs00269932_m1, MYOD1; Hs02330075_g1, HBG1/HBG2; Hs00361131_g1, IL1B; Hs01555410_m1, IL1R2; Hs01030384_m1, ACTB Hs99999903_m1. (Thermo Fisher Scientific).

## RESULTS

### Golden gate assembly of gRNA arrays with up to 10 gRNAs

To simplify the generation of gRNA arrays we utilized the golden gate cloning system [15]. We used type IIS restriction enzyme sites (*BsmBI*) and over hangs previously published and validated for robust golden gate assembly of TALEN DNA binding domains [18]. We then designed and assembled cassettes for oligonucleotide ligation of protospacer sequences using a different type IIS restriction enzyme (*BsaI*) flanking a stuffer sequence. In addition, we included a Csy4 ribonuclease target sequence directly upstream of the target gRNA sequences such that after golden gate assembly each gRNA is directly flanked by the Csy4 target sequence (**Fig. 1a**). Next, we designed a gRNA array acceptor plasmid (pACPT) containing a LacZ gene, for blue/white colony selection after golden gate assembly, flanked by appropriate *BsmBI* sites and an upstream U6 pol II promoter to drive expression of assembled gRNA arrays (**Fig. 1a**). We also included a terminal Csy4 target sequence, such that the last gRNA is free of additional sequence when processed, and a poly T termination sequence. In order to produce a highly modular system for rapid and efficient cloning of the U6 driven gRNA array cassettes, attL1/2 sequences were included in pACPT for Gateway cloning (**Fig. 1a**). We have termed this set of plasmids for oligonucleotide ligation pGG1-10 and the acceptor plasmids pACPT1-10 (**Fig. 1b**). An example of the plasmids required for golden gate assembly of a 4 gRNA array and the structure of the final expression plasmid are shown in **Fig. 1c**.

**Fig. 1.**
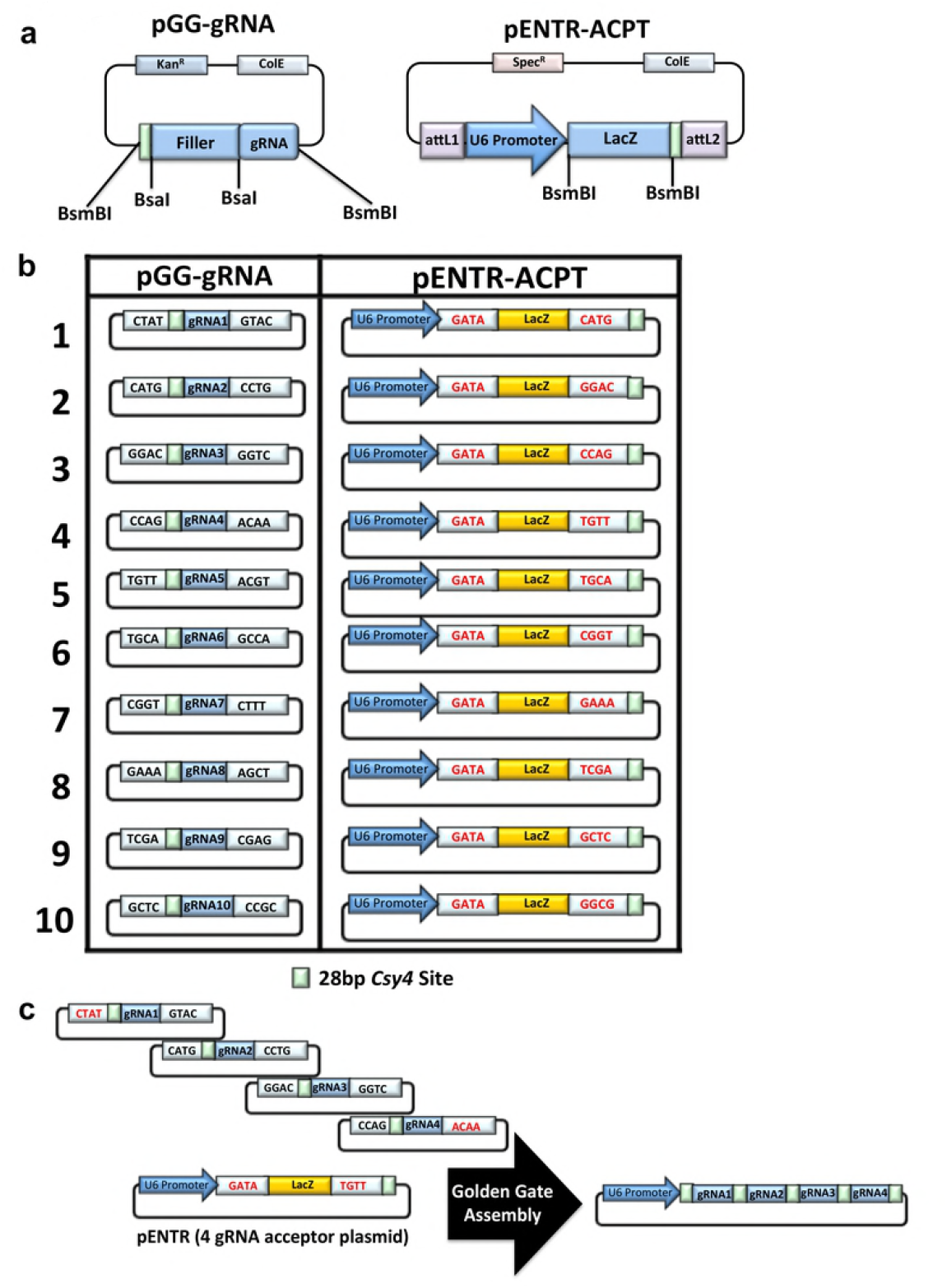
Golden gate assembly of gRNA arrays. (**a**) Diagram of the base pGG (Left) and pENTR-ACPT (Right) plasmids highlighting the type IIS restriction enzymes used for protospacer oligonucleotide ligation (BsaI) and golden gate assembly (BsmBI). In addition, the pGG cassette contains a filler sequence that is removed upon enzyme digestion and a 5’ Csy4 site (light green) for array processing once assembled and expressed. A terminal Csy4 site was included in the pENTR-ACPT cassette to remove additional plasmid sequence from the terminal gRNA when expressed and a LacZ gene that is removed upon golden gate assembly to allow for blue/white colony selection. (b) Diagram of the final 10 pGG and 10 pENTR-ACPT plasmids for assembly of arrays containing 1-10 gRNAs. The gateway attL1/2 sites of pENTR-ACPT plasmids have been left out for simplicity. (c) Example of the plasmids required to assemble an array of 4 gRNAs.

In order to validate our golden gate assembly system we ligated oligonucleotides encoding protospacer sequences targeting 10 genes we previously validated for CRISPR/Cas9 mediated DSB induction with an average editing frequency of ~22% and a range of 10-35% (1:*GOSR1*, 2:*PPP2R2A*, 3:*CNTFR*, 4:*DMD*, 5:*ZBTB10*, 6:*KAT7*, 7:*SPPL3*, 8:*CCM2*, 9:*PRDX1*, 10:*TRIP12;* **S1 Fig. 1**). We then performed golden gate assembly of arrays containing 3, 5, 7, or 10 of these gRNAs and validated the resultant plasmids by Sanger sequencing (**S2 Fig. 2**). As with the previously described golden gate assembly system, we observed sufficient white bacterial colonies upon X-gal staining and all white colonies sequenced as expected. These data demonstrate that we have developed a highly functional platform for the generation of gRNA arrays using our golden gate cloning platform.

**Fig. 2.**
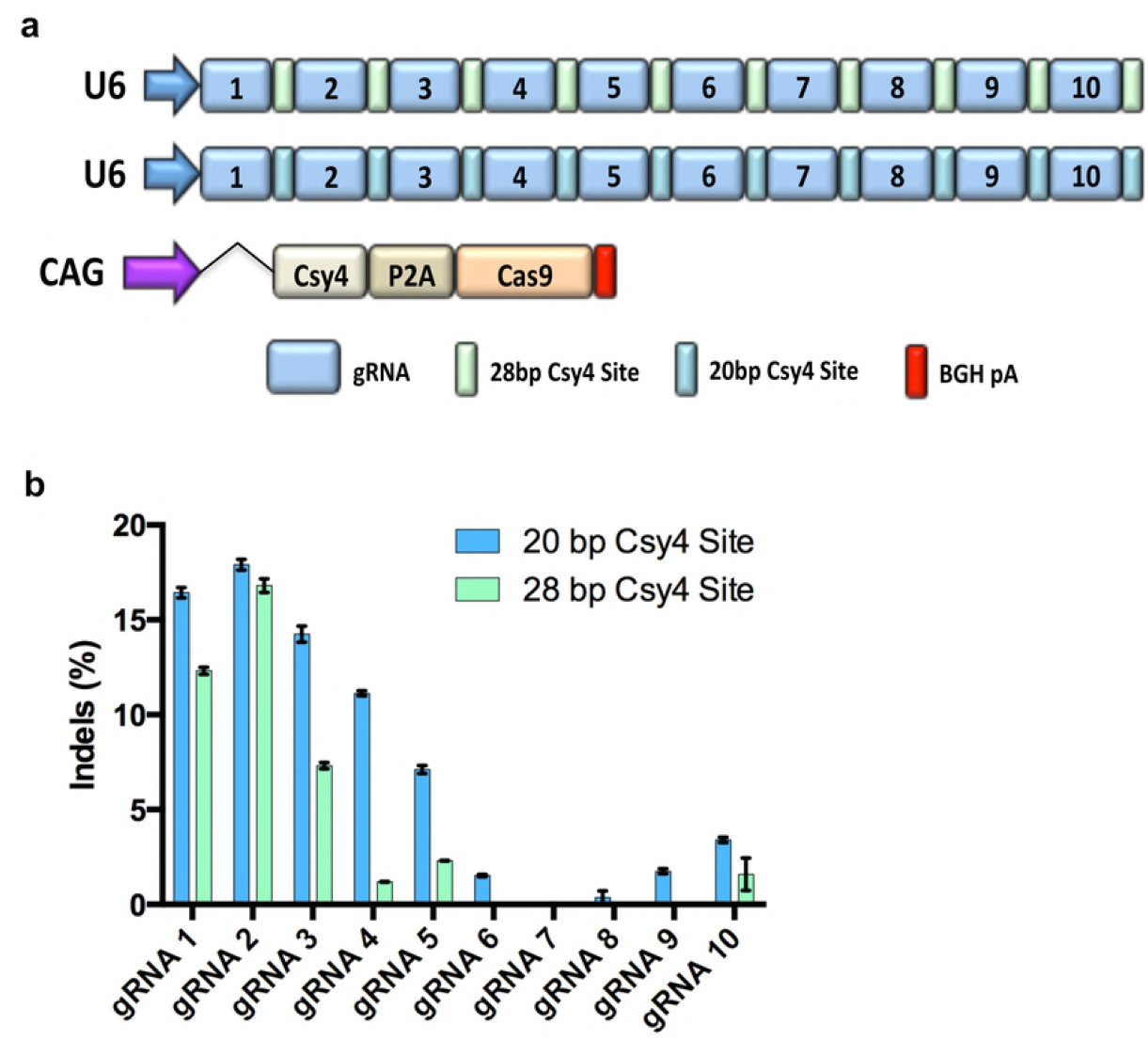
Gene editing frequency of pol III driven 10 gRNA array. (a) Diagram depicting the plasmid vectors transfected into HEK293T cells to induce targeted DSBs using a pol III driven 10 gRNA array combined with Cas9 and Csy4. (b) Results of CRISPR/Cas9 editing at each of the 10 gRNA target sites when using gRNA arrays with a 20 or 28bp Csy4 target sequence. Mutation frequencies were assessed by Surveyor Nuclease assay with means of triplicate measurements shown. P2A: ribosomal skip sequence; BGH pA: bovine growth hormone polyadenylation signal; CAG: strong mammalian promoter comprised of cytomegalovirus (CMV) early enhancer element, the first exon and intron of chicken beta-actin gene, and the splice acceptor of the rabbit beta-globin gene.

### Assessment of gene editing frequencies using gRNA arrays

In order to validate the functionality of our assembled gRNA arrays, we transfected HEK293T cells with gRNA arrays containing 3, 5, 7, or 10 gRNAs and a plasmid expressing Cas9 alone or Cas9 linked via a P2A element to the human codon optimized Csy4 ribonuclease [19] (**Fig. 2a**). Notably, we observed negligible editing without expression of *Csy4* to process the array into individual gRNAs, confirming the necessity of *Csy4* for array processing (**S3 Fig**). The results of nuclease activity for the 10 gRNA array transfected with Cas9-P2A-Csy4 demonstrated detectable rates of editing with the first four gRNAs in the array and then the editing diminished to nearly undetectable levels at gRNA 7 and 8, but editing was again observed with the 9^th^ and 10^th^ gRNA (**Fig. 2b**).

Interestingly, two previous reports using 2 or 4 gRNA arrays differed in the length of flanked *Csy4* target sites used, 20bp or 28bp [20, 21]. The larger 28bp *Csy4* sequence contains a ‘handle’ region of 8bp that may be important for *Csy4* processing^19^. However, there is no clear evidence demonstrating which flanking *Csy4* target sequence is superior for optimal *Csy4* processing. Thus, we generated an additional set of pGG1-10 and pACPT plasmids harboring the 28bp Csy4 target site and again assembled the 10 gRNA array expressed via the U6 pol III promoter. The gRNA array containing the 20bp Csy4 site produced slightly higher levels of gene editing at all 10 target sites, indicating the 20bp Csy4 site may be more efficiently cleaved by *Csy4* than the 28bp sequence (**Fig. 2b**). However, we still observed low editing over all and nearly undetectable editing at gRNA 7 and 8. This phenomenon was even less prominent with shorter 3 and 5 gRNA arrays (**S4 Fig**). These data demonstrate that golden gate assembled gRNA arrays with an optimal 20bp Csy4 site can induce detectable gene editing at up to 10 target sites, but the editing is highly reduced after the initial 4 gRNAs in the array when using larger arrays.

### Pol II promoter driven gRNA arrays produce enhanced gene editing frequencies

While a 20bp Csy4 cleavage site leads to slightly higher gene editing frequencies than a 28bp site, the gene editing frequencies are lower than individual U6-gRNA editing overall. We hypothesized that the use of the canonical U6 driven expression of the highly repetitive array may produce a very unstable transcript due to the lack of a polyA tract and G cap associated with pol III driven genes. Thus, in order to stabilize the gRNA array transcript, we removed the U6 promoter from the pACPT1-10 plasmids (**S5 Fig. 5**) and again assembled an array of 10 gRNAs that were subsequently cloned into a vector containing the strong pol II CMV promoter with a poly adenylation sequence (**Fig. 3a**). The array was transfected into HEK293T and demonstrated improved gene editing frequencies overall, with ~10%-21% gene editing across all gRNA targets, albeit still at lower frequencies than individual U6-gRNA editing (**Fig. 3b**, green line vs blue dots). We then tested the 10 gRNA array expressed from the very strong intron containing pol II CAG promoter with a poly adenylation sequence (**Fig. 3a**). Surprisingly, we found the rates of gene editing using the CAG promoter were significantly higher than the individual U6-gRNA editing, with only 3 gRNA target sites having slightly lower editing when expressed in CAG driven array (**Fig. 3b**, purple line, *p* < 0.001). We further tested the effect of the promoter driving the expression of arrays containing 3, or 5 gRNAs and again observed optimal gene editing using the CAG promoter (**S4 Fig**). These results demonstrate gRNA arrays expressed from strong pol II promoters with polyadenylation sequences enhance gene editing frequencies to levels as high or higher than observed with individual standard U6-gRNA plasmids.

**Fig. 3.**
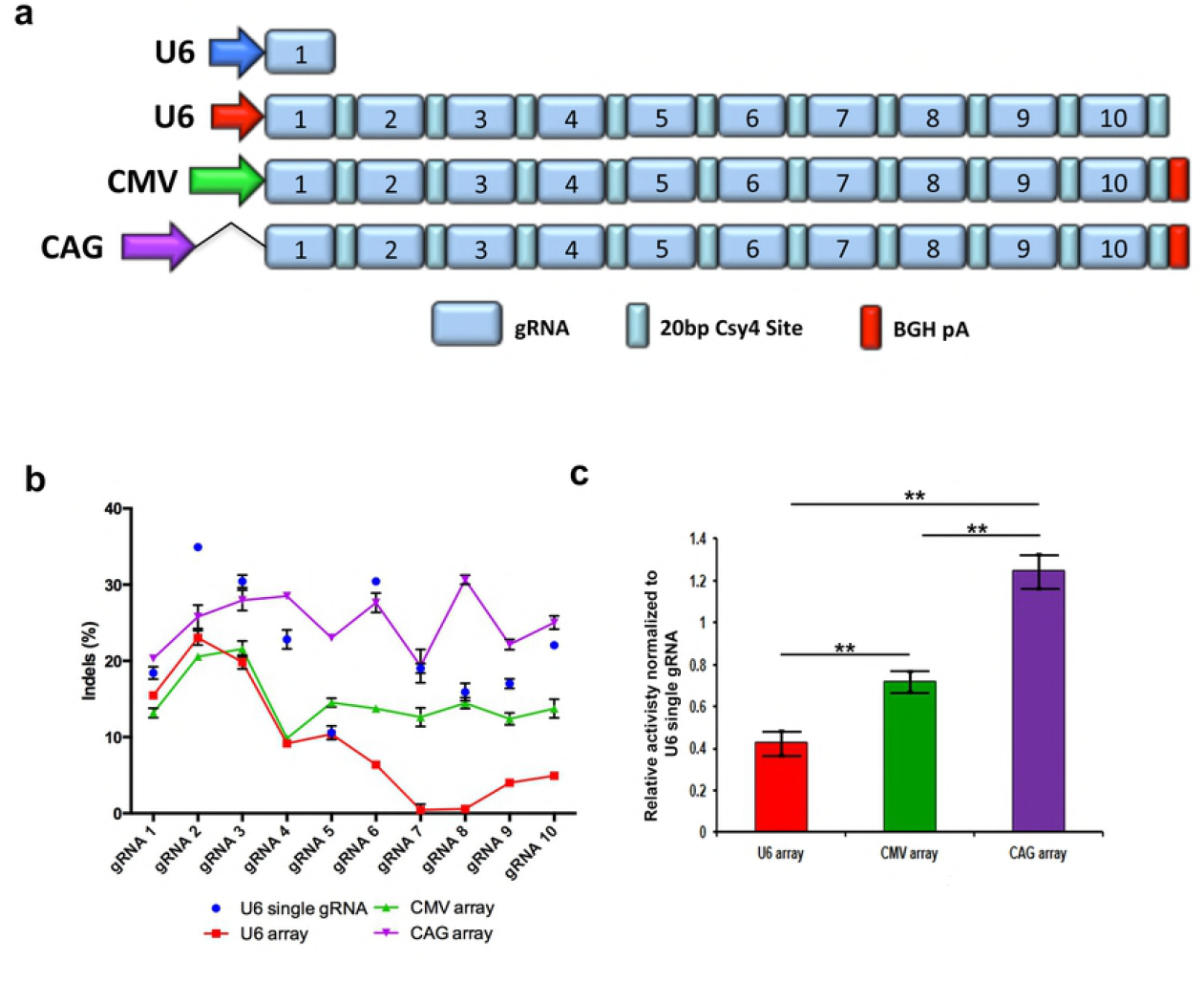
Comparison of gene editing frequency of pol II and pol III driven gRNA arrays. (a) Diagram depicting the plasmid vectors transfected into HEK293T cells containing pol II or pol III promoters driving transcription of a 10 gRNA array. (b) Graph depicting the gene editing frequency of each gRNA when expressed as individual gRNAs transcribed from the standard U6 pol III promoter (blue dots) or in a single 10 gRNA array transcribed from the standard U6 pol III promoter (red line), CMV promoter with BGH polyadenylation signal (green line), and CAG promoter with BGH polyadenylation signal (purple line) assessed 3 days post transfection. (c) Bar graph depicting the average gene editing frequency of the 10 gRNA array expressed from each promoter normalized to the editing frequency of each individual gRNA transcribed from the standard U6 pol III promoter. Mutation frequencies were assessed by Surveyor Nuclease assay with means of triplicate measurements shown. P2A: ribosomal skip sequence; BGH pA: bovine growth hormone polyadenylation signal; CMV: cytomegalovirus; CAG: strong mammalian promoter comprised of CMV early enhancer element, the first exon and the first intron of chicken beta-actin gene, and the splice acceptor of the rabbit beta-globin gene. \*\**p* < 0.001, *p*-values are from two-way ANOVA with Tukey’s post hoc test. Error bars, SEM

### gRNA arrays produce significantly higher gene editing over multiplexed plasmid delivery

One of the motivations for the development of a robust gRNA array system is to avoid the toxicity and low transfection efficiency associated with delivery of numerous U6-gRNA containing plasmids. Thus, we wanted to test our gRNA array plasmids against with standard multiplexed plasmid delivery. We first attempted to deliver 10 individual U6-gRNA plasmids along with a Cas9 expressing plasmid using 250 ng of each plasmid as we did with our gRNA array and Csy4/Cas9 vector (**Fig. 4a**). However, this produced substantial toxicity and death of nearly all cells transfected, likely due to the use of a large amount of plasmid. Next, we reduced the amount of each of the 10 individual U6-gRNA plasmids to 25 ng along with Cas9 to equal the same amount (in micrograms) as our gRNA array and Csy4/Cas9 vector. This resulted in no obvious toxicity and analysis of gene editing efficiency at all 10 target sites was significantly higher using our gRNA array approach (**Fig. 4b**). The average editing efficiency for 10 individual U6-gRNA plasmids were significantly lower (8.2%) compared to with the gRNA array (25.0%). These data demonstrate the gRNA array system is a superior approach to the use of numerous multiplexed U6-gRNA plasmids.

**Fig. 4.**
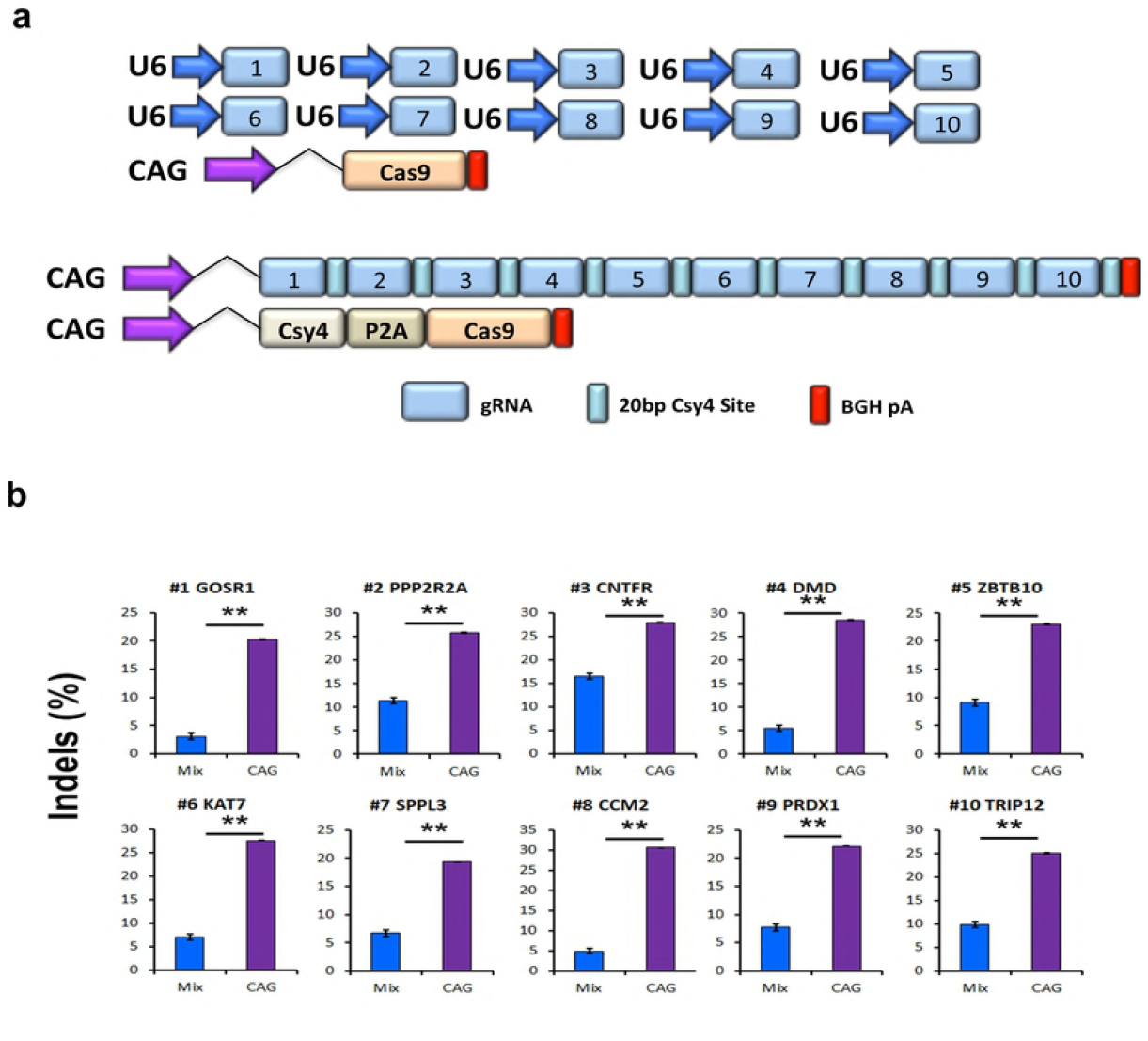
Enhanced multiplex editing using gRNA arrays. (a) Diagram depicting the plasmid vectors transfected into HEK293T cells to compare gene editing by multiplexing 10 standard U6- gRNA plasmids and a 10 gRNA array. (b) Bar graph depicting the gene editing frequency at each of 10 gRNA target sites 3 days post transfection using multiplexed individual U6-gRNA plasmids or 10 gRNA array encoding the same gRNAs. Mutation frequencies were assessed by Surveyor Nuclease assay with means of triplicate measurements shown. P2A: ribosomal skip sequence; BGH pA: bovine growth hormone polyadenylation signal; CAG: strong mammalian promoter comprised of CMV early enhancer element, the first exon and the first intron of chicken beta- actin gene, and the splice acceptor of the rabbit beta-globin gene. \*\**p* < 0.00, Student’s *t* test. Error bars, s.d.

### gRNA arrays can work with Pol II promoters in multiplex synergistic activation mediator (SAM) activation

In order to allow for multiplex gene activation using gRNA arrays, we generated a set of pGG1-10 vectors with gRNAs containing 2 MS2 binding sites (**S6 Fig**). These gRNAs, driven by a strong pol II promoter, are compatible with the highly effective SAM activation system [3]. We used this system to generate a gRNA activation array containing 5 previously validated gRNAs used for gene activation [3] (**Fig.** *5***a**). We then transfected HEK293T cells with individual U6-MS2-gRNAs, dCas9-VP64 and MS2:p65:HSF1 plasmids to assess standard gene activation with the SAM system using individual U6 gRNAs (**Fig. 5b**). These activation results were then compared with gene activation in cells transfected with the MS2 gRNA array, Csy4/dCas9-VP64 and MS2:HSF1:p65 plasmids. We observed robust levels of gene activation and, therefore these results demonstrate that MS2 gRNA arrays are amenable to multiplex gene activation.

**Fig. 5.**
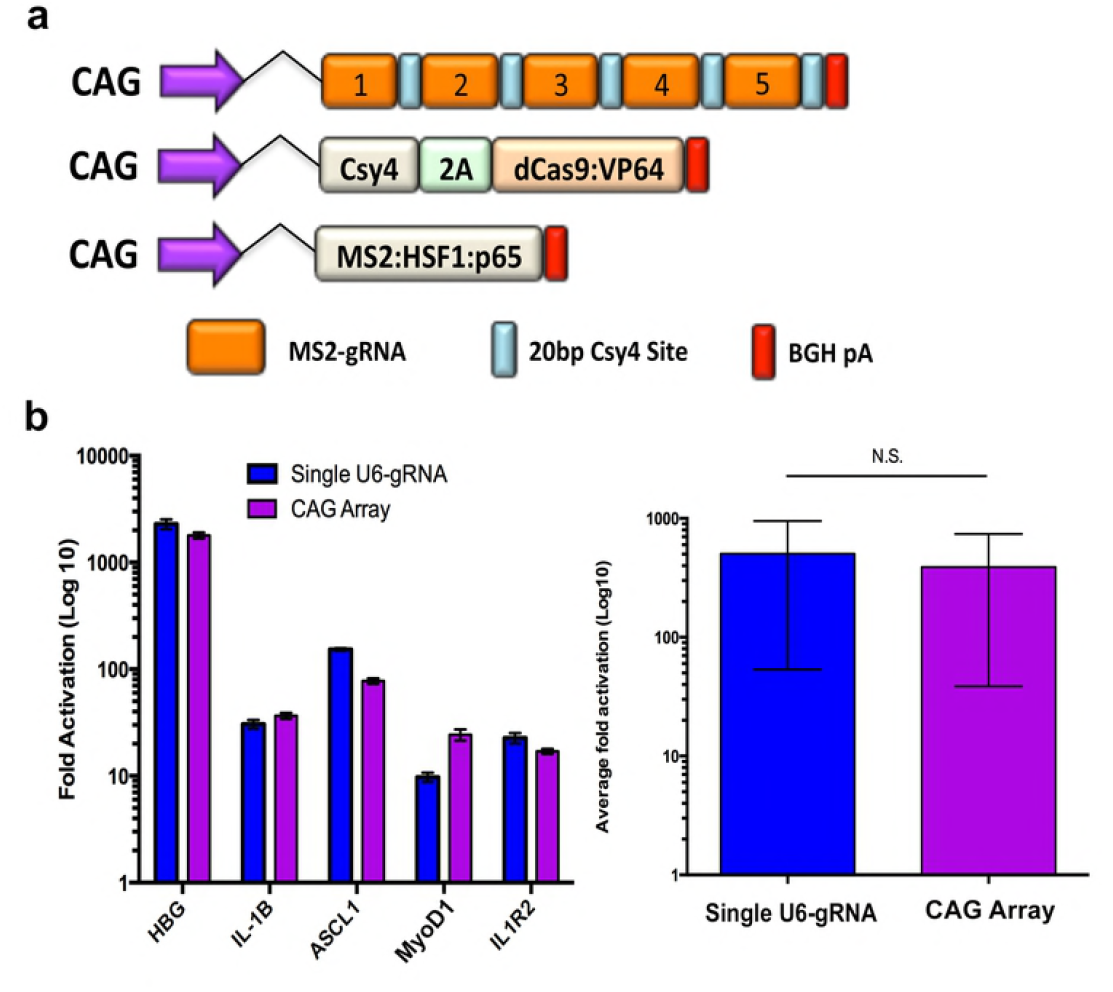
Multiplexed gene activation using the SAM system with gRNA arrays. (a) Diagram depicting the elements encoded in plasmids used for multiplex gene activation using the SAM system combined with gRNA arrays containing MS2 sequences. (b) RT-PCR results of gene activation at 5 gRNA target sites 3 days post transfection using individual U6-gRNAs or a gRNA array containing all 5 gRNAs (left). Average gene activation using either approach is also shown, demonstrating no difference in gene activation using single U6-gRNA plasmids or gRNA arrays (right).

## DISCUSSION

We report the development and methods for assembly of CRISRP/Cas9 gRNA arrays capable of expressing up to 10 gRNAs from a single promoter. These gRNA arrays are effectively processed by *Csy4* ribonuclease and high rates of gene editing can be detected at all gRNA target sites when expressed from the strong pol II CAG promoter containing a polyadenylation sequence. Moreover, gene editing frequencies are higher when using pol II driven gRNA arrays compared to the individual standard U6-gRNA plasmids, especially when multiplexing numerous U6-gRNA plasmids.

One of the more unexpected results of our study was that the use of the pol II CAG promoter optimally expressed the gRNA array, in that it resulted in higher gene editing frequencies compared to individual U6-gRNAs plasmids, which is the most commonly used format of the CRISPR/Cas9 system. This is unexpected as the U6 promoter has been shown to be highly efficient at transcription of gRNAs with nearly a log fold higher expression compared to CMV for instance [19]. Although the CAG promoter may simply produce larger amounts of transcript compared to the standard U6 promoter, another potential reason for enhanced editing may be due to the use the Csy4 enzyme. Perhaps Csy4 protects the gRNA from degradation that normally occurs from endogenous non-specific RNases, providing a larger window of time for Cas9 to bind the gRNA and induce targeted DSBs. Alternatively, Csy4 may directly interact with Cas9 to enhance gRNA loading after gRNA array processing. Future studies investigating the mechanism of Csy4 array processing and conceivable interaction with Cas9 will be required to identify any potential mechanism of enhancement of the system.

Since its discovery the CRISPR/Cas9 system has been rapidly adopted for numerous applications due to its ease of use, specificity, and the seemingly limitless ability to maintain function when fused to various protein domains. However, some of the applications, such as chromosome labeling and genetic circuits, require the use of numerous gRNAs at one time. For instance, Chen et al. found that labeling unique genomic regions using the dCas9-EGFP fusion required 26-36 gRNAs to obtain a signal to noise ratio allowing the region to be visible by microscopy [22]. Chen et al. achieved stable expression of gRNAs by packaging each gRNA in individual lentiviral vectors and transducing target cells with the pooled lentiviral vectors. There have also been a number of publications using the CRISPR/Cas9 system to generate genetic circuits, though only a few gRNAs are typically utilized [19, 22, 23] and at maximum 7 gRNAs were actually used in the experiments [8, 21]. As this field progresses to generate more sophisticated genetic circuits the requirement to express more gRNAs will also likely increase.

Although we developed a gRNA array platform for *sp*Cas9 using standard and MS2 containing chimeric gRNA backbones, similar platforms can likely be developed for other CRISPR orthologs (such as *Neisseria meningitidis* Cas9 and *Staphylococcus aureus* Cas9), other modified gRNA backbones, and other CRISPR systems (such as *Cpf1*). Moreover, it is also possible to generate golden gate assembly libraries to mix and match various gRNA backbones to use multiple orthologs simultaneously. For instance, *Sp* dCas9-VP64 for gene activation combined with *Sa* dCas9-KRAB for gene repression using a gRNA array containing both *Sp* and *Sa* specific gRNA backbones. The use of *Cpf1* for enhanced multiplex genome engineering is especially intriguing based on recent work demonstrating that Cpf1 has both DNase and RNase activity. Fonfara et al. established that Cpf1 uses its DNase activity to induce sequence specific DSBs and its RNase function to process the transcribed CRISPR arrays into individual gRNAs [21]. Thus, by using *Cpf1* it may be possible to deliver just one protein to carry out the analogous function of Cas9 and Csy4 in the system described here.

Finally, a very exciting aspect of the gRNA array technology is the ability to use *in vitro* transcribed RNA encoding the gRNA array. This approach may be especially desirable for multiplexed editing of primary human lymphocytes, such as T cells. It is well documented that plasmid DNA is highly toxic to primary lymphocytes [24] and thus the use of IVT gRNA arrays may allow for highly efficient multiplex gene editing of primary human cell types for research and therapy. Indeed, we were able to generate and test IVT gRNA array in primary human T cells combined with mRNA encoding Csy4 and Cas9. However, we observed no detectable gene editing using this approach in primary human T cells (data not shown). This negative result may be due to a timing issue as the gRNA array is perhaps largely degraded by the time active Cas9 and Csy4 protein become abundant in the cells. This issue has been recently described using single IVT gRNAs in T cells [7]. Perhaps the use of purified Cas9 and Csy4 protein combined with IVT gRNA arrays will allow for highly multiplexed editing of primary cell types. An alternative approach would be to deliver the entire system with AAV6, which has been recently shown to be highly effective at delivering transient transgene expression in both primary human T cells and CD34^+^ hematopoietic stem cells[16, 25].

The use of gRNA arrays for highly multiplexed genome engineering using the CRISPR system will likely help expand and enhance the ability to perform multiplexed genome engineering for basic research and therapies.

## ACKNOWLEDGEMENT

B.S.M., N.K.W., and M.K. designed the research. M.K., M.S. performed the QRT-PCR analysis. M.T.W, W.S.L, S.L., K.H., B.R.W, N.K.W. and M.K. constructed all vectors. N.K.W., M.K., and S.L. performed all transfections. N.K.W. and W.S.L performed all Surveyor nuclease assays. B.S.M. wrote the article.

